# Green tea and Spirulina extracts inhibit SARS, MERS, and SARS-2 spike pseudotyped virus entry in vitro

**DOI:** 10.1101/2020.06.20.162701

**Authors:** Jeswin Joseph, Karthika T, Ariya Ajay, V.R. Akshay Das, V. Stalin Raj

## Abstract

Coronaviruses (CoVs) infect a wide range of animals and birds. Their tropism is primarily determined by the ability of the spike (S) protein to bind to a host cell surface receptor. The rapid outbreak of emerging novel coronavirus, SARS-CoV 2 in China inculcates the need for the development of hasty and effective intervention strategies. Medicinal plants and natural compounds have been traditionally used to treat viral infections. Here, we generated VSV based pseudotyped viruses (pvs) of SARS-, MERS-, and SARS-2 CoVs to screen entry inhibitors from natural products. In the first series of experiments, we demonstrated that pseudotyped viruses specifically bind on their receptors and enter into the cells. SARS and MERS polyclonal antibodies neutralize SARSpv and SARS-2pv, and MERSpv respectively. Incubation of soluble ACE2 inhibited entry of SARS and SARS-2 pvs but not MERSpv. In addition, expression of ACE2 and DPP4 in non-permissive BHK21 cells enabled infection by SARSpv, SARS-2pv, and MERSpv respectively. Next, we showed the antiviral properties of known enveloped virus entry inhibitors, Spirulina and Green tea extracts against CoVpvs. SARSpv, MERSpv, and SARS-2pv entry were blocked with higher efficiency when preincubated with either green tea or spirulina extracts. Green tea provided a better inhibitory effect than the spirulina extracts by binding to the S1 domain of spike and blocking the interaction of spike with its receptor. Further studies are required to understand the exact mechanism of viral inhibition. In summary, we demonstrate that pseudotyped virus is an ideal tool for screening viral entry inhibitors. Moreover, spirulina and green tea could be promising antiviral agents against emerging viruses.

## Introduction

The past three decades have witnessed a tremendous increase in the number of highly pathogenic emerging viruses [1] which have caused serious global threats initiating diverse approaches for understanding the biology of the virus and for robust vaccine and therapeutic development. Coronaviruses are notable examples of zoonotic emerging viruses which can infect a diverse range of mammals and birds including humans [2]. The major outbreak of human coronaviruses occurred in 2003 wherein Severe Acute Respiratory Syndrome (SARS) CoV emerged from Chinese palm civet and caused an epidemic in China with more than 8000 cases including 774 fatal cases [3]. With the help of public health authorities, the outbreak was effectively contained within six months. Ten years later, another CoV termed Middle East respiratory syndrome (MERS) CoV emerged from dromedary camel to human and is continuing to cause outbreaks in the Middle East region [4]. To date, 2519 cases have been reported with mortality rates of 34% [5]. There is no approved therapy or vaccine available for this virus yet. By the end of 2019, another pandemic coronavirus named Severe acute respiratory syndrome coronavirus 2 (SARS-CoV-2) emerged in Wuhan, China [6] though unknown intermediate host. Till now, more than 8,000,000 cases and 450,000 deaths have been reported worldwide [7]. Likewise, novel or re-emerging viral outbreaks may occur in the near future as well. Therefore, effective antiviral and therapeutic strategies are required to control the ongoing or future outbreaks.

Coronavirus spike protein is a promising target for the development of antiviral compounds because their cell tropism is primarily determined by the ability of the spike (S) protein to bind to a host cell surface receptor and can block the virus entry at the early stage of infection. But drug discovery for highly pathogenic viruses like SARS, MERS, Ebola, Lassa, or SARS-2 are challenging due to the requirement for a BSL-3/BSL-4 laboratory containment facility. However, very limited facilities are available, especially in developing countries. Pseudotyped viruses provide a substitute model in which the native envelope protein of a nonpathogenic BSL-2 virus (vesicular stomatitis virus) replaced with an envelope glycoprotein of a highly pathogenic virus like SARS, MERS, Ebola, or SARS-2 [8]. These viruses mimic a normal virus but are noninfectious in nature. Moreover, they are replication-incompetent with a single round of infection and hence can be used to do research in the routine BSL-2 laboratories. Pseudotyped viruses have been used in diagnostics, vaccines, and high-throughput screening of entry inhibitors for several BSL-3/BSL-4 level pathogens [8].

Natural products such as plants, algae, and seaweeds have always been implicated in multiple fields of biology for their antibacterial, antiviral, and antifungal properties. Several medicinal plants have been traditionally used to treat viral infections [9] and have been demonstrated with their ability of their inhibitory effects (Table 1) on the replication or entry of several viruses like herpes simplex virus (HSV) type 2, hepatitis B (HBV), influenza virus [10]–[13]and also other emerging viral pathogens such as poxvirus and SARS [14]. Curcumin acts as an antiviral agent against several viruses such as Parainfluenza virus type 3, vesicular stomatitis virus (VSV), HSV, etc. [15]. Diammonium glycyrrhizin the main component of licorice root extract shown to have an inhibitory effect on pseudorabies virus (PrV) [16]. Neem and Tulsi leaf extracts are potent antiviral agents against influenza virus [17], [18]. Moreover, the extract of green tea, another natural compound, inhibits human immunodeficiency virus (HIV), zika virus, influenza virus, and hepatitis C virus (HCV) [19].

**Table 1.** List of antiviral activity of naturally available plant and seaweed extracts

| Source | Target viruses | Mode of action | References |
| --- | --- | --- | --- |
| Olive leaves | Viral hemorrhagic septicemia rhabdovirus (VHSV), Hepatitis B virus (HBV) and Human Immunodeficiency Virus (HIV) | Cell to cell fusion, Secretion of HBV antigens, Fusion and integrase inhibition | [39]–[41] |
| Licorice Root | SARS-CoV, Human Respiratory Syncytial Virus (HRSV) HIV | Viral entry and replication | [42]–[44] |
| Gigartina Skottsbergii | Human Herpesvirus 1 and 2 (HHV 1 and 2) Herpes simplex virus type 1&2 (HSV-1 and 2) | Cell surface attachment and replication | [38] |
| Spirulina | HIV-1<br>Herpes simplex virus type 1 and 2 (HSV-1 and 2)<br>Vesicular stomatitis virus (VSV)<br>Influenza virus | Viral Entry, Replication, Cell membrane fusion<br>Hemagglutination, | [21], [45],<br>[46] |
| Green Tea | HIV-1<br>HSV-1,2<br>Influenza virus)<br>Hepatitis B and C virus (HBV, HCV)<br>Dengue virus (DENV)<br>Zika virus (ZIKV) | Reverse transcriptase inhibition<br>Cell surface attachment<br>Neuraminidase activity inhibition<br>Interference in replication and transcription<br>Viral entry<br>Viral particle destruction | [19], [26],<br>[47], [48] |

Spirulina is a commercially available dietary supplement which has been recorded for its diverse properties. It is a free-floating cyanobacterium, which has 70% protein content and is rich in phenolic acids, essential fatty acids, sulfated polysaccharides, and vitamin B12 [20]. Extracts of Spirulina have been shown to have antiviral activity against multiple enveloped viruses including influenza virus, HSV, adenovirus, etc.[21] Apart from crude plant extracts, several specific compounds from Green tea, catechins, such as epicatechin (EC), epigallocatechin (EGC), epicatechin gallate (ECG), and epigallocatechin-3-gallate (EGCG) have been found to have antiviral and anticarcinogenic properties [22] (Supplementary table 1). EGCG is a major component and active catechin of green tea and has several bio modulatory effects such as anti-allergic, anti-inflammatory, anti-tumor, antioxidative, and antiviral properties. EGCG, has been reported to inhibit HCV, HIV, HSV type 1 and 2, enterovirus 71, influenza A, and other viruses [10], [23]–[27]. For HIV, EGCG binds to the CD4 molecule at high affinity and inhibits HIV gp120 binding to human CD4+ T cells [28]. Moreover, the EGCG-induced inhibition was observed in a broad spectrum of HIV-1 subtypes as well. EGCG acts through direct inactivation of the virus particle, inhibition of the protease adenain, and intracellular growth in vitro [29]. Sulfated polysaccharides and Spirulan like compounds are the major components of spirulina extract, which inhibits the entry of several viruses including HSV-1, hepatitis A, human cytomegalovirus (HCMV), VSV, and HIV [13]. This clearly shows the importance of antiviral properties of natural compounds, and hence could be promising therapeutic agents for emerging viral diseases including SARS-CoV 2. In this study, we show the biological applications of pseudotyped coronaviruses and the antiviral activity of Spirulina and green tea extracts using the developed pseudotyped coronaviruses.

## Materials and Methods

### Cells

Vero E6 and HEK293T cells were grown in DMEM (Lonza, 12-604F) supplemented with 10% FBS (MP, 29101) and 1% penicillin/streptomycin. BHK21 cells were maintained in EMEM (Lonza, 12-611F) supplemented with 10% FBS and 1% penicillin/streptomycin (Lonza, 17-602E). Huh7 cells were grown in RPMI 1640 medium (Lonza, 12-115F) with 10% FCS and 1% penicillin/streptomycin. The cells were grown at 37°C in a CO2 incubator.

### Plasmids

Angiotensin converting enzyme 2 (ACE 2) gene (2.4kb) was PCR amplified from cDNA of Vero E6 cells and the resulted amplicon was inserted between KpnI/XbaI restriction sites in PCDNA3.1+ expression vector (pCDNA-ACE2). Dipeptidyl peptidase 4 (DPP4)(2.3kb) was amplified from cDNA of Huh7 cells, as described previously [30], cloned between BamHI/NotI in PCDNA3.1+ expression vector (pCDNA-DPP4). A synthetic construct of the full-length spike of SARS-CoV-2 (aa residues 1-1273) was commercially synthesized from Genewiz, UK. The coding sequence of spike S1 domain of SARS-2(aa residues 1-683), SARS (1-676), and MERS (1-747) were amplified from synthetic spike construct and cDNA of SARS- and MERS-CoV respectively. Amplicons were fused C-terminally with Fc domain of human immunoglobulin (IgG) (aa residues 1-231) inserted via SacI /XhoI restriction sites in pCAGGS expression vector (pCAGGS-SARS2-S1-Fc, pCAGGS-SARS-S1-Fc, pCAGGS-MERS-S1-Fc). For the production of soluble ACE2, gene sequence encoding amino acid residues 1-635 was PCR amplified from pcDNA-ACE2 plasmid introduced between N terminal CD5 signal sequence and C Terminal 6X HIS-tag in pCAGGS-CD5 plasmid by Kpn1/Xho1 (pCAGGS-sACE2).

### Production of pseudotyped coronaviruses

Full-length spike genes of SARS (aa residues, SARS-2(1-1254) and MERS CoV lacking the endoplasmic retention signal sequence were cloned into pCAGGS expression system (pCAGGS-SARS2-S, pCAGGS-SARS-S, and pCAGGS-MERS-S) and were transfected using Polyethylenimine (PEI) (Polysciences, USA) at 1:3 ratio in HEK293T cells. After 24 h post-transfection, cells were infected with VSVΔG/GFP pseudovirus, incubated for 1 hour, and cells were washed thrice with PBS and added with infection medium DMEM with 1%FCS. The cell supernatant was harvested after 24h post-infection (pi), cell debris was removed by centrifugation at 3000rpm for 10 minutes and aliquots were stored at −80°C. For titration assay, a day before the experiment Vero E6 and Huh7 cells were seeded in a 96 well plate at 2×10^4^ cells per well and 100μl of serially diluted (1:10) pseudoviruses were added to the cells and incubated for 1h. After incubation, cells were washed and replaced with fresh DMEM with 1% FBS and were incubated at 37°C in a CO2 incubator for 24h. GFP positive cells were counted and were normalized to VSVΔG –GFP background values. The lowest dilution of the pseudovirus with GFP positive cells was calculated and viral titer was calculated as infection units /ml as mentioned previously [31], [32].

### Transmission Electron Microscopy (TEM)

Pseudoviruses were concentrated using Amicon 3kDa cut off columns (Merck, UFC800324). Tem micro liters of glutaraldehyde (2.5%) fixed samples were drop casted on formvar carbon-coated copper grids and negatively stained with 1% Uranyl acetate alternative solution for 1 minute (TED PELLA, Cat. No-19485). The grids were washed twice with distilled water to remove the excess stain and followed by imaging on Tecnai, FEI Transmission electron microscope at 120 kV.

### Production of recombinant proteins

Ten micrograms of plasmids encoding either spike S1-Fc or sACE2 (pCAGGS-SARS2-S1-Fc, pCAGGS-SARS-S1-Fc, pCAGGS-MERS-S1-Fc, and pCAGGS-sACE2) were independently transfected using PEI in HEK293T cells. After 12h post-transfection, cells were replaced with fresh, freestyle expression media, and the cells were incubated for 5 days at 37°C in a CO2 incubator. Recombinant Fc fused S1 proteins containing supernatants were harvested and cell debris was removed by centrifugation for 5 minutes at 1500rpm. S1 -Fc proteins were purified using Protein A-Sepharose beads (GE Healthcare Life sciences) following the manufacturer’s instructions. Proteins were eluted from the column using 0.5M glacial acetic acid pH 3 and neutralized with 3M Tris-Hcl PH 8.8. Quality and quantity of the purified proteins were analyzed by nanodrop, BCA, and SDS-PAGE. Further confirmed by western blotting using Goat-anti human IgG conjugated with HRP (Bethyl, A80-119P). Soluble ACE 2 was purified using Nickel -NTA agarose beads (Qiagen Cat.No-1018244), eluted with 200mM imidazole, and was analyzed by SDS PAGE and western blot using Goat anti-human ACE 2 polyclonal antibody (R&D, Cat.No-AF933).

### sACE2 blocking assay

sACE2 (5μg/ml PBS) was preincubated with SARS2pv, SARSpv, and MERSpv at 37°C for 1h and then the mixture was directly added on Vero E6 cells for 1h. After incubation, the cells were replaced with fresh DMEM with 1% FCS and antibiotics and incubated for 24h at 37°C in a CO2 incubator. Next day, GFP positive cells were counted and calculated as the relative percentage of infection.

### Pseudovirus neutralization assay

SARS and MERS CoV polyclonal antibodies (gifted by Bart Haagmans, EMC, The Netherlands). SARS polyclonal antibodies were preincubated with SARSpv and SARS-2pv and MERS polyclonal with MERSpv (1:100 dilution) at 37°C for 1h. Then the antibody virus mixture was directly added on Vero E6 cells. 1hpi, infection medium was replaced with fresh DMEM with 1% FCS and was incubated at 37°C in a CO2 incubator for 24h. GFP positive cells were counted to calculate the antibody neutralization of the pseudoviruses.

### Surface expression of ACE2 and DPP4

BHK21 or HEK293T cells were transfected with **either** pCDNA-ACE2 or pCDNA-DPP4 using PEI at 1:3 ratio. After 6h post-transfection, the medium was replaced with complete DMEM with 10% FCS and 1% antibiotics. Surface expression was analyzed by incubation of either Goat anti-ACE2 or Goat anti-DPP4 (1:250) primary antibody and secondary antibody with rabbit anti-goat conjugated with Alexa Fluor 594 (Immunotag-ITIF59418). The positive cells were visualized using Leica TCS SP5 II inverted confocal microscope 63x oil objective.

### Cold water extraction of Spirulina (*Arthrospira platensis*) and Green Tea

Spirulina and green tea are available in routine organic/medical stores. Spirulina was purchased as capsules or powder and green tea was purchased as dried leaves. Extracts were prepared as described elsewhere [33]. Briefly, Spirulina and green tea were powdered using mortar and pestle and the fine powder was weighed at 20mg/ml concentration and were mixed in sterile distilled water and vortexed for 5 min. Next, the mixtures were freeze-thawed and the supernatants were collected by centrifugation. Supernatants were filter sterilized using a 0.22μm membrane filter and stored at 4 °C until further use.

### Screening of antivirals against pseudotyped coronaviruses

In the first series of experiments we incubated the cells with either Spirulina, green tea extracts or DMEM. After 30 minutes incubation, cells were replaced with 100microlitre of all the four pseudotyped viruses. Parallelly, we incubated all the fours pvs as well with the extracts or DMEM. After 30 minutes of incubation, the mixtures were directly added to confluent Vero E6 cells and incubated for 1h. Then the cells were washed and added with fresh DMEM 1% FCS and antibiotics. Next day, GFP positive cells were counted and the percentage of infection was calculated.

### Spike and receptor interaction studies by S1-Fc binding analysis

HEK293T cells were transfected with either pCDNA-ACE2 or pCDNA-DPP4 plasmids. 24h post-transfection, spike S1-Fc proteins (5μg/ml) of SARS, SARS-2 and MERS CoV and the cells were independently treated with Spirulina and green tea extract for 90 min at 37°C. In the first set of experiments, untreated spike S1-Fc proteins were added to pretreated cells followed by binding assay at 4°C. In the next set of experiments, the preincubated extract - spike S1-Fc protein mixture was added to the untreated cells and were processed for binding assay. In both the experiments, following 1h of spike S1-Fc incubation at 4°C, the cells were stained with Goat anti-human IgG conjugated with FITC (1:300 dilution; A80-119F, Bethyl) subsequently counterstained with DAPI (Sigma Aldrich Cat no D9542-10MG). Confocal images were acquired using Zeiss LSM 880 confocal laser-scanning microscope with an objective at 63x oil. Imaging parameters were kept the same for control versus treated samples. The images were processed using the Zeiss ZEN blue software.

### Statistical analysis

Unpaired student’s t-test was performed and only p values of 0.05 or lower were considered statistically significant. For all statistical analyses, the GraphPad Prism 5 was used.

## Results

Emerging viruses are highly pathogenic and require sophisticated biocontainment facilities. Research on these pathogenic viruses like SARS, MERS, or SARS-CoV-2 is challenging due to the limited availability of BSL-3/BSL-4 laboratory facilities. Pseudotyped viruses are an alternative that can mimic a live virus, but replication-deficient and can be used for research work in the routine BSL-2 laboratory. For the generation of coronavirus pseudotypes, full-length spike gene of SARS, MERS, and SARS-2 lacking the endoplasmic retention signal sequence were amplified from cDNA (SARS and MERS) or the codon-optimized synthetic construct (SARS-CoV-2) and amplicons were cloned into pCAGGS expression plasmid. The plasmid constructs were confirmed by sequencing and restriction analysis and were transfected in HEK293T cells, and protein expression was confirmed by immunofluorescent staining (data not shown). In order to produce VSVΔG/GFP pseudotyped coronaviruses, cells were transiently expressed with the respective plasmids encoding the spike glycoprotein of SARS, MERS, or SARS-CoV-2 in HEK293T cells and VSVΔG/GFP pseudovirus was infected 24 hours post-transfection. Pseudo typed coronaviruses were harvested from the precleared cell supernatant and titrated in Vero E6 or HuH 7 cells, and the GFP positive cells (Fig 1a) were counted and calculated the infection units per milliliter (IU/ml) The titer of the SARSpv was 1.2×105 IU/ml, MERSpv 1.4×104 IU/ml and VSVpv 1×106.4 IU/ml whereas SARS CoV-2 yielded a less titer 5×103 IU/ml (Fig. 1b). The presence of intact CoVpv particles in the supernatant was analyzed independently by Transmission electron microscopy (TEM). Intact bullet-shaped virus particles of 200 nm size were observed in each grid (Fig. 1c).

**Figure 1:**
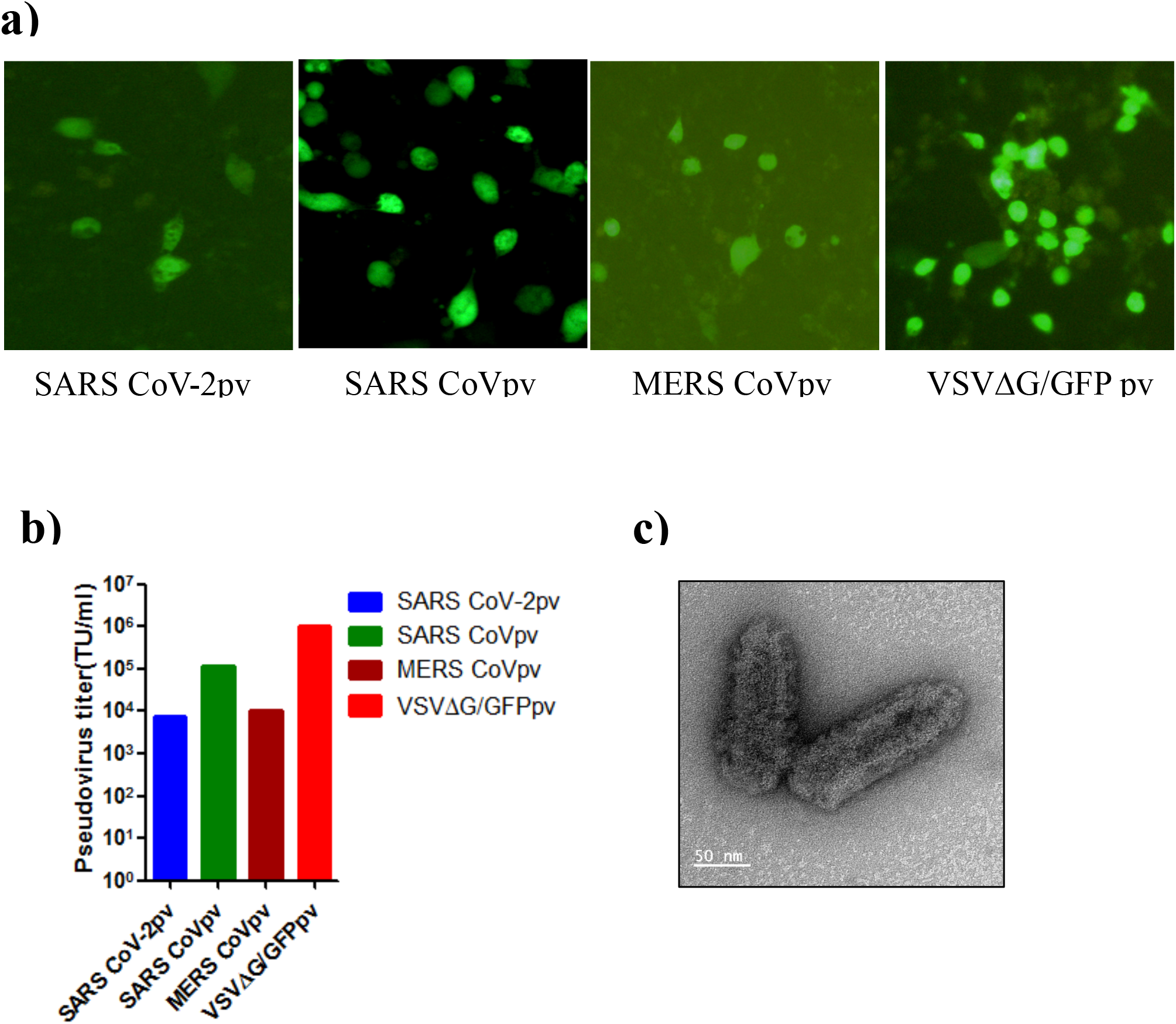
Generation of VSV pseudotyped coronaviruses. **a)** Infection of SARSpv, SARS-2pv, MERSpv and VSVpv in Vero E6 cells. GFP expression shows the infected cells. A single GFP positive cell corresponds to a single pv infection (200x zoomed image). **b)** Serially diluted pseudotyped viruses were titrated on Vero E6 and Huh7 cells. The lowest dilution showing GFP positive cells were calculated and viral titer is represented as Infection units /ml (IU/ml). **c)**. Presence of intact pseudotyped viruses were aanalsyed by negative staining and transmission electron microscopy which depicted pseudovirus particles of size 200nm. Representative image of pseudotyped SARS CoVpv.

Proper maturation of the VSV pseudotyped coronavirus particles are critical for cellular entry and infection. Next, we checked whether the polyclonal antibodies raised against MERS or SARS CoVs binds and neutralizes the pseudotyped CoVs. SARS-CoV specific polyclonal antibodies neutralizes SARSpv and also cross neutralizes SARS-2pv but not MERSpv (Fig 2a) whereas, MERS-CoV polyclonal specifically neutralizes MERSpv but not SARSpv or SARS-2pv (Fig. 2b), suggesting that antibodies specifically bind on the receptor binding S1 domain of the CoVpvs and neutralize the virus infection. It is well known that SARS and SARS-CoV-2 use human ACE2 as the entry receptor to initiate an infection cycle [33]–[35]. To confirm that the CoVpv entry is mediated by its specific receptor, we produced a recombinant soluble form of Angiotensin-converting enzyme 2 (sACE2) in HEK293T cells and purified using Ni-NTA affinity column. A 90 kDa band was observed in SDS PAGE and was confirmed by western blotting using ACE2 specific antibody (Fig. 2c). Incubation of sACE2 with CoVpvs blocked the entry of SARSpv and SARS-2pv but did not affect MERSpv infection (Fig 2d).

**Figure 2:**
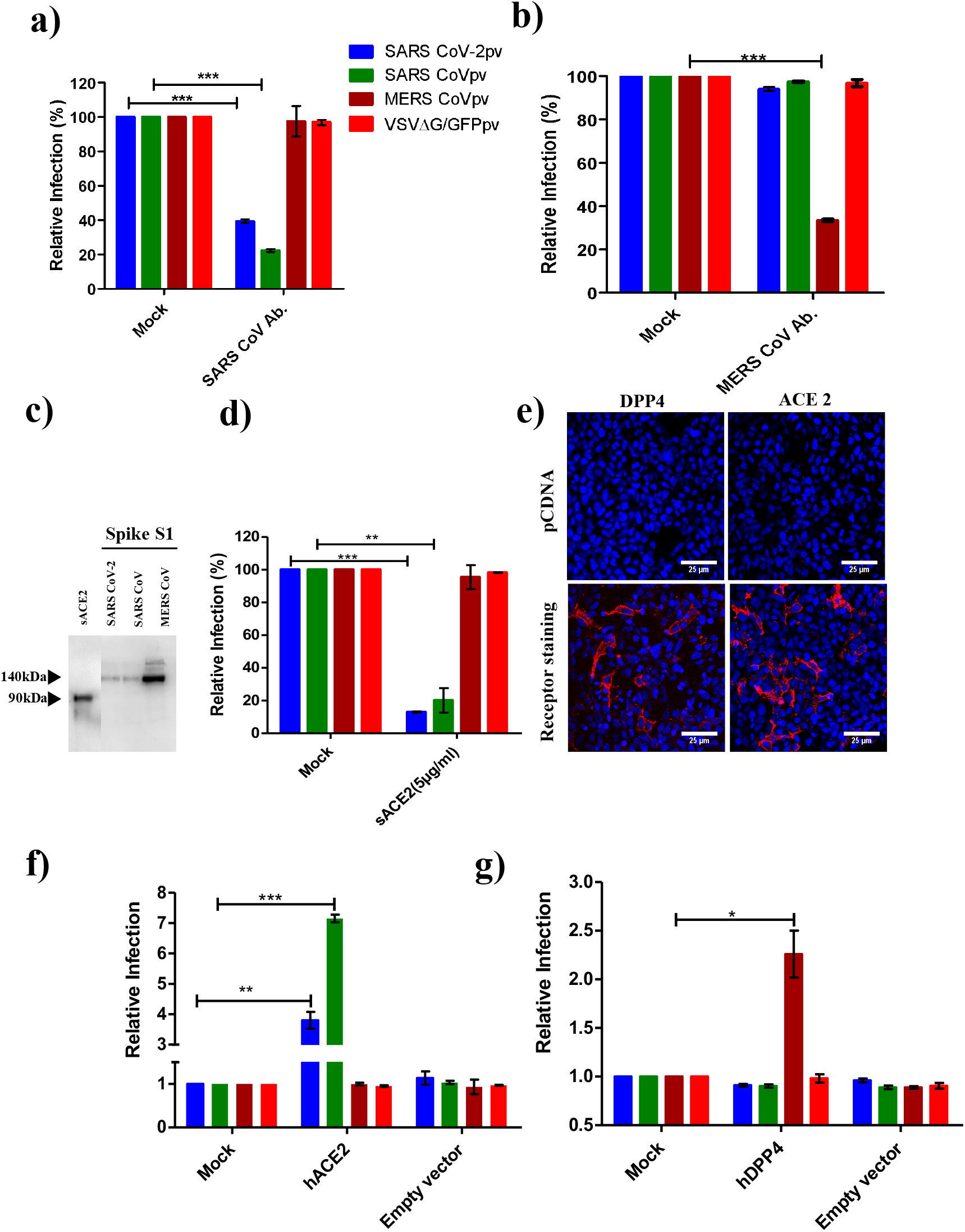
Antibody neutralization and receptor specificity of the developed pseudoviruses. Maturation of the developed pseudoviruses were assessed neutralization assay with **a**) SARS CoV polyclonal antibody. Significant reduction in infection was observed in SARS and SARS −2pv **b)** MERS CoV polyclonal antibodies and significant reduction was observed only in MERSpv but not in others. **c)** 6X HIS-tagged recombinant sACE2 was expressed in Hek293T cells and was purified by Ni-NTA based IMAC purification. The purified protein was confirmed by western blotting using anti-ACE2 antibodies to detect a band of ≈90KDa. Similarly, SARS-, SARS-2- and MERS-CoV spike S1 fused with Human Fc, the recombinant protein was produced and purified with protein A sepharose beads. Approximately, 140kDa protein was purified and was confirmed by western blotting using Goat anti human IgG HRP conjugated antibody. **d)** In order to validate the specific interaction of the pseudotyped particles on mammalian cells, soluble ACE2 (sACE2) was incubated with pvs followed by pv infection on Vero E6 cells**. e)** Surface expression of either ACE2 or DPP4 on BHK21 cells were analyzed by immunostaining using Goat anti-ACE2 or goat Anti-DPP4 antibody. Non permissive BHK21 cells were transiently expressed with **f)** ACE2 or **g)** DPP4 followed by pvs infection. SARSpv and SARS-2pv were susceptible in ACE2 expressed cells and MERSpv was susceptible in DPP4 expressed cells. Data shown is relative normalized infection and error bar represents SD. Unpaired t test was performed for statistical analysis. P values, P<0.05 =* P<0.01=** and P<0.001=***

Further, to confirm the receptor-mediated entry of the pseudotyped coronaviruses, we cloned the complete ACE2 or MERS-CoV receptor dipeptidyl peptidase 4 (DPP4) and was transiently expressed on the non-susceptible BHK21 cells. The surface expression of ACE2 and DPP4 was confirmed by immunostaining using ACE2 and DPP4 polyclonal antibodies (Fig 2e). Next, we expressed the S1 domain of different spike proteins fused to the Fc domain of human IgG (S1-hFc) in HEK293T cells, yielding recombinant proteins of approximately 140 kDa, which was confirmed by western blotting (Fig 2c) and immunostaining (Supplementary Fig 1). Incubation of recombinant SARS-, SARS-2 -S1-hFc fusion proteins enables binding on hACE2 whereas MERS-S1-hFc bind on the hDPP4 transfected cells and no binding was observed in empty plasmid transfected cells (Fig 2e). Subsequently, ACE2 transiently expressed on the surface of non-susceptible BHK21 cells enable SARS- and SARS-2 pvs infection, whereas MERSpv or VSVpv did not (Fig. 2f). Similarly, MERS CoVpv infection was observed in hDPP4 expressed BHK21 cells, whereas SARS-, SARS-2-, and VSV-pvs infections were similar to empty plasmid transfected cells (Fig 2g). These data confirm that the developed pseudotyped CoVs use its specific receptor to enter into the host cells.

Targeting viral entry through interaction with cell receptors is a preliminary goal for the development of antivirals. Medicinal plants and natural compounds have been previously reported to have antiviral activity against a diverse range of viruses [9], [36]. Among these, several natural compounds are known to inhibit viral entry by binding to the viral glycoprotein or host cell surface receptor and block the interaction of the virus and its receptor, thereby preventing the entry into the host cells. Spirulina and Green tea extracts have been reported to have the ability to inhibit the viral entry of enveloped viruses, including HCV, Influenza, HIV, VSV, etc. [21], [23], [26], [27], [37]. Here, we screened the antiviral activity of Spirulina and Green tea extracts against pseudotyped coronaviruses. First, to know whether crude extracts inhibit CoVpvs, we treated Vero E6 cells as well as pseudotyped viruses (SARSpv, SARS −2pv, and MERSpv), including a VSVpv positive control with 0.2mg/ml of cold water extracts (CWE) of Spirulina and green tea.

Interestingly, a significant reduction in infection was observed in all pvs pretreated with the extracts (Fig 3b, d) but not in any of the preincubated cells (Fig 3a, c). The results suggest that CWE of Spirulina and Green tea inhibits pseudotyped coronavirus entry by attachment to pseudovirus envelope protein rather than to the cell surface.

**Figure3:**
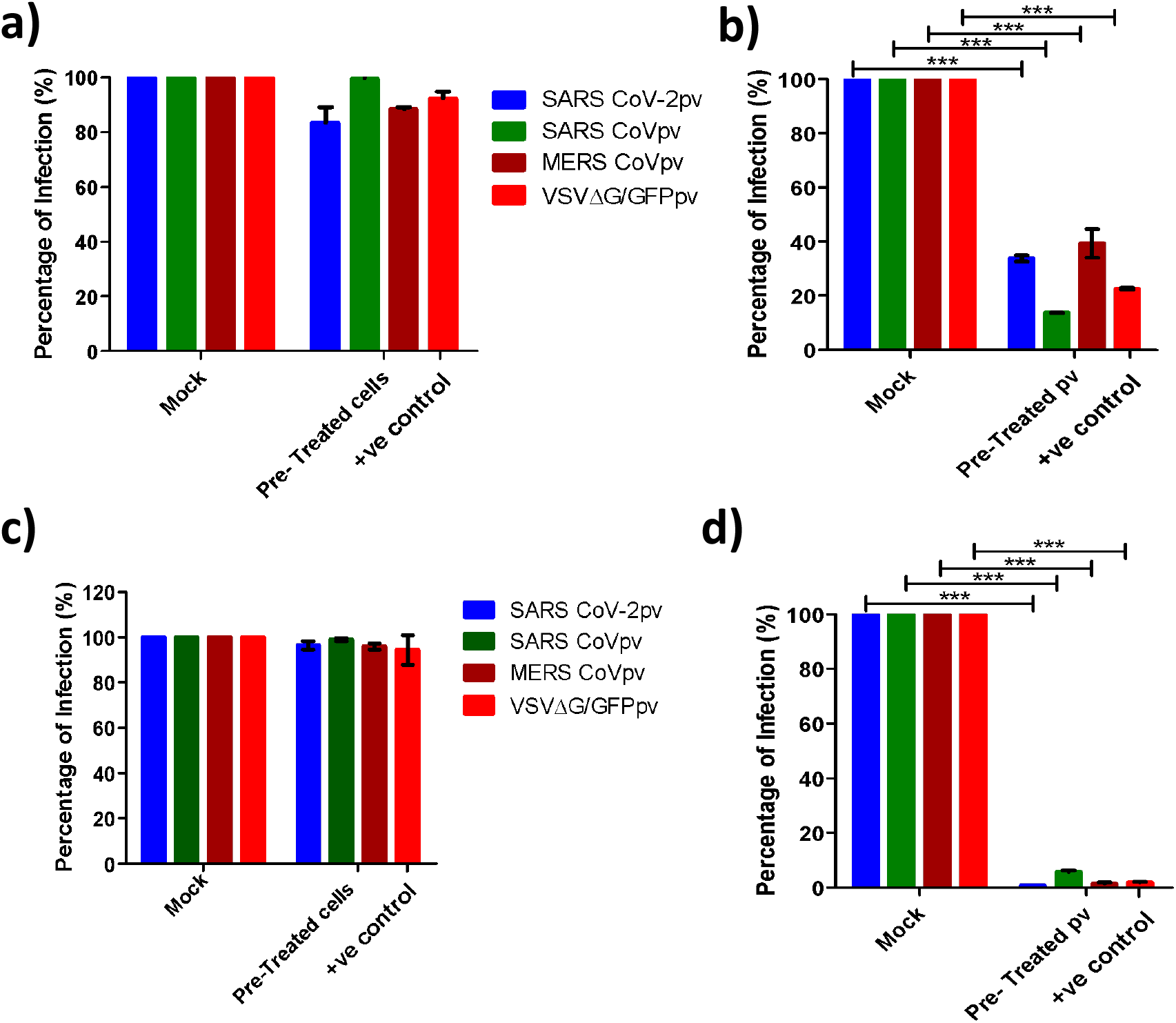
The cold water extract of Spirulina and Green Tea inhibits viral entry in a broad range of coronaviruses *in vitro*. **a and c) Monolayer** of Vero E6 cells were incubated either Spirulina or green tea extract (0.2mg/ml) followed by pvs infection. **b** and **d)** pseudoviruses were pre-incubated with Spirulina extract or green tea extract (0.2mg/ml) and then the mixture was directly incubated on Vero E6 cells. Data represent the mean ± SD of an experiment with duplicates. The experiment was repeated twice. Statistical significance was determined by unpaired t test. P value, P<0.001 =***

Next, to confirm that the observed inhibitory activity of the extracts was not due to proteolytic cleavage of the spike glycoprotein, we incubated the spike S1-hFc protein with the extracts for 90 minutes and then the proteins were analyzed by western blotting. No protein degradation was observed in any of the proteins in comparison to input (Supplementary 2). Next, we performed a concentration-dependent inhibition assay for all pvs. We incubated pvs with varying concentrations ranging from 0.1mg/ml −0.8mg/ml of Spirulina and green tea extracts. All three CoVpvs were confirmed to be inhibited by CWE of Spirulina and Green tea with higher efficiency. 90% inhibition was observed in Spirulina treated SARSpv, SARS-2pv, MERSpv and VSVpv infection at 0.6mg/ml,0.5mg/ml,0.8mg/ml and 0.6 mg/ml respectively (Fig.4a)whereas green tea treated pseudotyped viruses were inhibited at 0.1mgml, 0.12mg/ml, 0.1mg/ml and 0.25mg/ml respectively (Fig.4b). Most importantly, Green tea CWE showed more inhibitory activity on pseudotyped coronaviruses in comparison to Spirulina extract.

**Figure 4:**
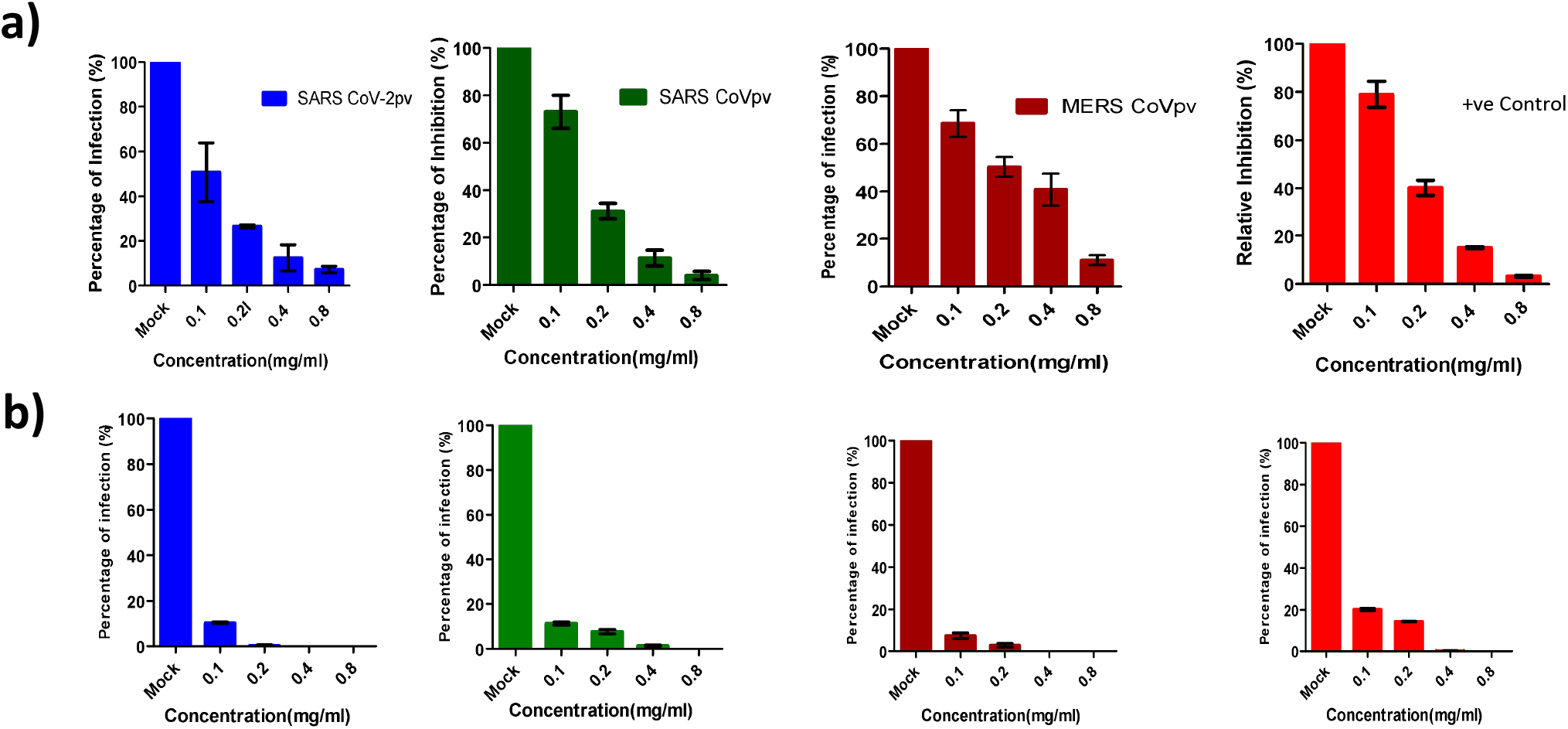
Concentration dependent inhibition of peusotyped viruses by *Spirulina* and green tea extracts. Different concentrations (0.1mg/ml −0/8 mg/ml) of Spirulina and green tea extracts were tested for the antiviral activity. **a)** represents Spirulina treated pseudotyped coronaviruses and **b)** depicts green tea treatment followed by infection assay. Blue bar represents - SARS-2pv, green-SARSpv, brown -MERSpv and red -VSVpv. Data represent the mean ±SD of an experiment with duplicates. The experiment was repeated twice.

To understand whether the inhibitory mechanism of Spirulina and green tea extracts are mediated through inhibition of the interaction of spike and its receptor, we performed an S1 binding assay on HEK293T cells expressing either ACE2 or DPP4. Cells were preincubated with the extracts followed by the addition of S1 spike proteins at 4°C. Binding assay of spike (S1-hFc) proteins was analyzed by immunostaining followed by confocal microscopy. Similarly, we incubated the spike S1 proteins with the extracts and then the mixture was added on the ACE2 or DPP4 cells. Cells treated with either of the extracts did not inhibit the interaction of S1 and its receptor ACE2 or DPP4 (Fig.5a). In contrast, all three S1 proteins incubated with the extracts of green tea is unable to bind to its receptor whereas Spirulina treated S1 proteins did not inhibit S1 -ACE2 or DPP4 interaction (Fig.5b) suggesting that the Spirulina extract blocks the viral entry through an unknown mechanism. However, green tea extract binds to the S1 domain of the spike and prevents the spike receptor interaction. Further studies with live viruses are required to elucidate the exact mechanism of antiviral action.

**Figure 5:**
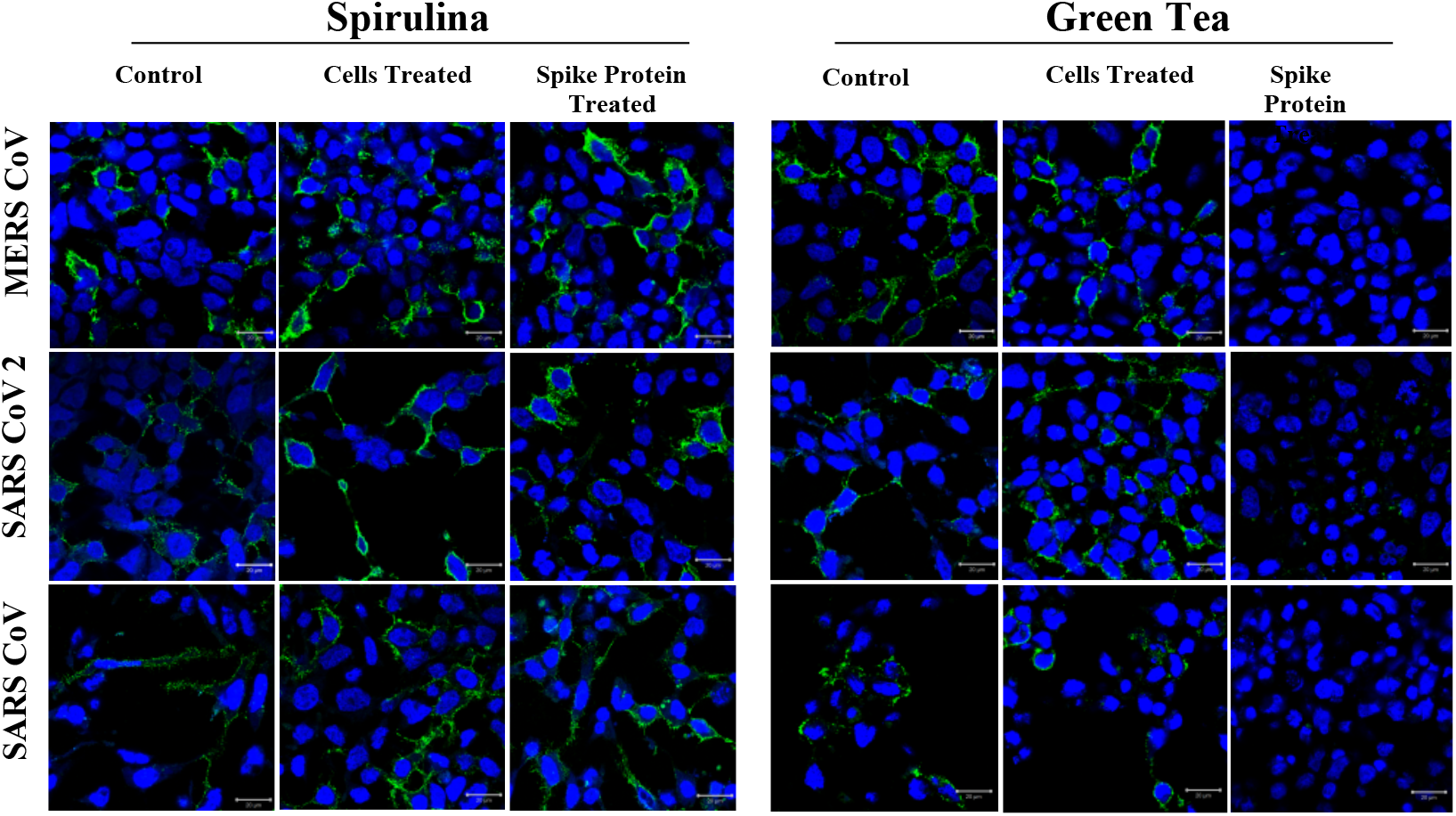
Pre-treatment of spike proteins with green tea extract inhibited S1-receptor interaction. **a, b**) DPP4 and ACE 2 plasmids were transiently transfected in HEK293T cells. 24 h post-transfection cells were independently incubated with PBS for 30 minutes followed by spike S1-fc protein (both left panels), spirulina and green tea extracts (0.4mg/ml) followed by S1-Fc protein (middle panels) and preincubated mixture of S1-Fc and Spirulina or green tea extract (right panels). Binding of S1-Fc was immunostained by Goat anti-human IgG conjugated with FITC and the binding of S1-Fc proteins were visualised by confocal microscopy. Green-IgG -FITC and Blue-DAPI. Scale bar 20μm.

## Discussion

The recent emergence of coronaviruses has caused serious global threats initiating diverse approaches for biocontainment of these pathogens. Most of these emerging viruses are highly pathogenic with limited therapeutic strategies and require BSL3/BSL4 facilities to handle these pathogens. The ongoing pandemic of SARS CoV-2 which has spread across the world, is a notable example that elucidates the need of diagnostics and robotic screening systems for identification of antiviral compounds. However, the limited high-level biosafety equipped laboratories particularly in developing countries inculcate the need for adapting to alternative approaches which can lead to the handling of the viruses in a BSL2 facility. Here we report the production of VSV pseudotyped virus particles for SARS CoV-2, SARS CoV, and MERS CoV. The developed pseudoviruses were susceptible for infection in Vero E6 cells and other cells. We obtained a high titer for SARS and MERS CoVpv whereas SARS CoV-2pv had a less titer which could be due to less packaging efficiency in HEK293T cells which was consistent with the previous report [31]. Pseudovirus production in Vero E6 cells [31] and expression of TMPRSS2 might enhance pseudovirus production and infection [33]. The developed pseudotyped viruses could be efficiently neutralized by specific polyclonal antibodies. SARS specific polyclonal neutralizes SARSpv and SARS-2pv as shown elsewhere [35] whereas MERS polyclonal antibodies neutralized MERSpv [36]. Therefore, pvs are useful tools for studying virus neutralization assay for highly pathogenic viruses in BSL2 facility [9]. Next, we showed that pseudoviruses specifically bind its receptor and enter into host cells. Previous studies report that pvs can be used to understand the entry of pathogenic viruses including coronaviruses [33].

The inhibition of viral entry is a key target for the development of antivirals. Medicinal plants and natural compounds have been reported to have a wide range of antiviral activities. Among these, Spirulina and Green tea extracts have been reported to inhibit attachment of enveloped viruses such as HCV, HIV, and Influenza virus [19], [21]. In our first study, we show that VSV is inhibited by the cold-water extract of Spirulina and Green tea. In the case of green tea consistent with previous studies, VSV entry is inhibited whereas cold water extracted Spirulina could inhibit the entry of VSV. However, hot water extract of Spirulina could not inhibit VSV entry [38] which might be due to the degradation of the major active compounds of Spirulina. but not hot water extract. Next, we tested the antiviral activity of Spirulina and Green tea cold water extract on pseudotyped coronaviruses. Treatment of cells with the extract did not inhibit viral entry whereas preincubation of pseudoviruses with the extracts inhibited viral entry on Vero E6 cells suggesting that the active compounds from Spirulina and green tea binds on the virus glycoprotein and block the virus entry. To further understand the exact inhibitory mechanism, we performed an S1 binding assay on DPP4 or ACE2 expressing cells using a mixture of S1 proteins preincubated with the Spirulina and green tea extracts. Interestingly, all three S1 proteins treated with green tea abrogate the interaction of spike with its receptor whereas Spirulina treated S1 proteins could bind to its cellular receptor. The observed results suggest that green tea binds to the spike S1 domain and inhibits the viral entry. However, we speculate that EGCG, EGC, ECG, or EC, the most active components in Green tea might be inhibiting the coronaviral entry not only binding exclusively to S1 but also other domains of the viral envelope glycoprotein. These compounds are reported to have antiviral entry inhibitory activity against different viruses [19]. On the other hand, the exact mechanism of action of Spirulina extract is not understood. The major components in Spirulina extract, sulfated polysaccharides and Spirulan like substances, might bind to enveloped glycoproteins of CoVs and play a role in the inhibition of coronavirus entry. Further research using the active compounds and live viruses is essential to understand the mechanism of action of these extracts. In summary, we have successfully developed pseudotyped viruses for SARS-SARS-2 and MERS CoV and demonstrated their applications including pseudovirus neutralization, virus entry and most importantly, screening for identification of antiviral compounds against viral entry. Moreover, natural products could be promising antiviral agents for emerging viruses.

## Acknowledgments

We thank Dr. Bart L. Haagmans and Mart M. Lamers, Erasmus MC, Netherlands for providing SARS and MERS CoV polyclonal antibodies and cDNA. VSVΔG/GFP pseudovirus was received from Yoshiharu Matsuura, Osaka University, Japan.We are grateful to the Central Instrumentation Facility provided by IISER TVM and Mr Alex Andrews for TEM assistance.

This study was approved by institute biosafety committee (IBSC) No. BT/BS/17/447/2011-PID

## Author’s contributions

V.S.R. designed and coordinated the study. J.J., K.T., A.A., and V.R.AD conducted the experiments. All authors contributed to the interpretations and conclusions presented. J.J, K.T and V.S.R. wrote the manuscript.

