## Supplementary Data for "Green tea and Spirulina extracts inhibit SARS, MERS, and SARS-2 spike pseudotyped virus entry in vitro"

**Supplementary file**

**Supplementary Table 1. List of antiviral activity of active compounds in spirulina and green tea extract.**

| Active compound | Target virus(es) | Mode of action | Reference(s) |
|---|---|---|---|
| | | **Spirulina Extract** | |
| Allophycocyanin | Enterovirus 71 | Reduced cytopathic effects and viral-induced apoptosis | [49] |
| Calcium spirulan | HIV-1, HSV-1, HCMV, Measles morbillivirus, Mumps virus, Influenza A virus, and HHV 8. | Inhibition of cytopathic effect, blocking viral attachment and penetration into host cells | [21], [37], [50] |
| Spirulan-like substances | HCMV, HHV-6 | Viral entry and intracellular pathway | [51] |
| Sulphoquinovosyl diacylglycerol | HSV-1 | Viral replication | [11] |
| Sulfated polysaccharides | HSV-1, Hepatitis A virus-type-MBB, HCMV, VSV, and HIV | Viral Entry | [13], [52] |
| | | **Green Tea Extract** | |
| Epigallocatechin (EGC) | Influenza virus, Adenovirus, and HIV-1 | Virus entry and replication | [27], [53] |
| Epicatechin gallate (ECG) | Influenza virus, Adenovirus, and HIV-1 | Virus entry, transcription and viral particle release | [27], [53], [54] |
| Epigallocatechin gallate (EGCG) | HIV-1, HCV, DENV, JEV, TBEV, ZIKV, Chikungunya virus | Viral attachment, integration, and replication | [23]–[26], [28], [29], [55]–[57] |
| | Influenza Virus | Alteration in the physical integrity of viral particles | [12], [27] |
| | Epstein Barr Virus (EBV) | Transcription of viral genes | [58] |
| | Adenovirus | Viral protein synthesis | [53] |
| Epicatechin (EC) | Influenza Virus | Viral entry | [27] |
| | Adenovirus | Viral protein synthesis | [53] |

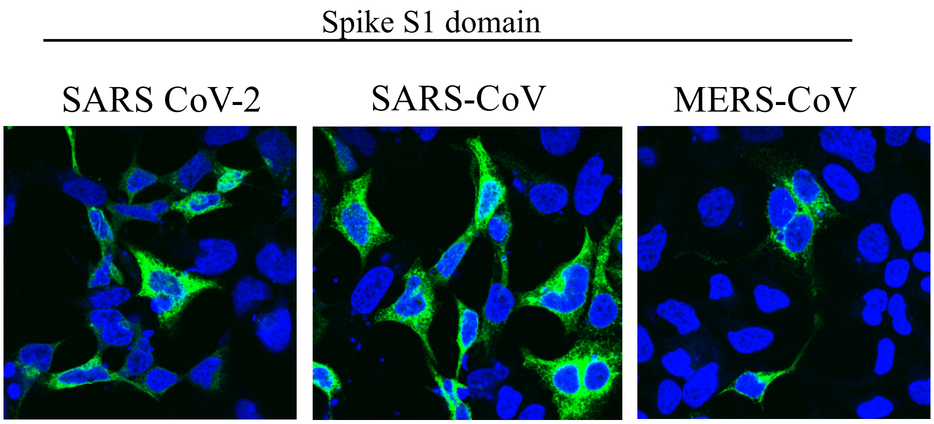


**Figure S1: Confirmation of recombinant spike s1-Fc tagged protein expression**. Generated constructs were transiently transfected in HEK 293T cell line. At 24 h of post transfection, proteins were stained with Goat anti human IgG FITC conjugated antibody (Green). Nuclei were stained with DAPI (blue)

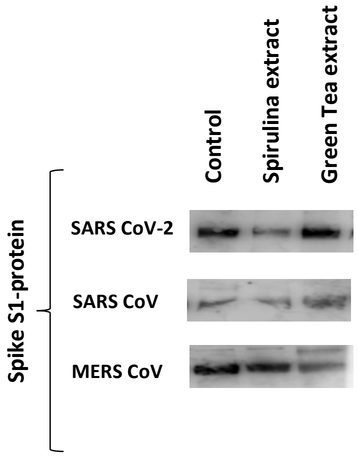


**Figure S2: Spirulina and Green tea extracts do not degrade envelope glycoproteins.** Spike S1 domain proteins (5µg/ml) of SARS-2, SARS and MERS CoV viruses were subjected for treatment with spirulina and green tea extracts (0.4 mg/ml) for 90 mins at 37°C and stained with Goat anti human IgG HRP conjugated antibody in western blot.
